# A Bacterial Surface Layer Protein Exploits Multi-step Crystallization for Rapid Self-assembly

**DOI:** 10.1101/665745

**Authors:** Jonathan Herrmann, Po-Nan Li, Fatemeh Jabbarpour, Anson C. K. Chan, Ivan Rajkovic, Tsutomu Matsui, Lucy Shapiro, John Smit, Thomas Weiss, Michael E. P. Murphy, Soichi Wakatsuki

## Abstract

Surface layers (S-layers) are crystalline protein coats surrounding microbial cells. S-layer proteins (SLPs) regulate their extracellular self-assembly by crystallizing when exposed to an environmental trigger. However, molecular mechanisms governing rapid protein crystallization *in vivo* or *in vitro* are largely unknown. Here, we demonstrate that the *C. crescentus* SLP readily crystallizes into sheets *in vitro* via a calcium-triggered multi-step assembly pathway. This pathway involves two domains serving distinct functions in assembly. The C-terminal crystallization domain forms the physiological 2D crystal lattice, but full-length protein crystallizes multiple orders of magnitude faster due to the N-terminal nucleation domain. Observing crystallization using time-resolved electron cryo-microscopy (Cryo-EM) reveals a crystalline intermediate wherein N-terminal nucleation domains exhibit motional dynamics with respect to rigid lattice-forming crystallization domains. Dynamic flexibility between the two domains rationalizes efficient S-layer crystal nucleation on the curved cellular surface. Rate enhancement of protein crystallization by a discrete nucleation domain may enable engineering of kinetically controllable self-assembling 2D macromolecular nanomaterials.

**Significance Statement:** Many microbes assemble a crystalline protein layer on their outer surface as an additional barrier and communication platform between the cell and its environment. Surface layer proteins efficiently crystallize to continuously coat the cell and this trait has been utilized to design functional macromolecular nanomaterials. Here, we report that rapid crystallization of a bacterial surface layer protein occurs through a multi-step pathway involving a crystalline intermediate. Upon calcium-binding, sequential changes occur in the structure and arrangement of the protein, which are captured by time-resolved small angle x-ray scattering and transmission electron cryo-microscopy. We demonstrate that a specific domain is responsible for enhancing the rate of self-assembly, unveiling possible evolutionary mechanisms to enhance the kinetics of 2D protein crystallization *in vivo*.

## Introduction

The structural determinants of macromolecular phase transitions are important to govern a wide variety of biological processes and disease states (1, 2). Microbial SLPs are highly expressed and ubiquitous proteins that self-assemble on the outside of bacteria and archaea to form crystalline protein coats (3–5). SLPs undergo a phase transition from aqueous to solid as part of their biological assembly and function (6, 7). RsaA, the 98 kDa SLP from *Caulobacter crescentus*, undergoes environmentally triggered self-assembly, forming sheets of crystalline protein with hexameric 22 nm repeats when exposed to calcium (7, 8). Recent work from our lab demonstrated that fast, efficient protein crystallization drives S-layer assembly *in vivo* (9). Therefore, we examined RsaA self-assembly *in vitro* using time-resolved small angle x-ray scattering (SAXS), circular dichroism spectroscopy (CD), and time-resolved Cryo-EM to determine the structural basis for S-layer assembly.

RsaA consists of two domains, both of which are necessary for successful S-layer assembly *in vivo* (Figure 1A) (10, 11). The N-terminal domain, here defined as RsaA_1-246_, is responsible for anchoring the protein to the cell surface via a non-covalent interaction with specific sugar moieties of the lipopolysaccharide (LPS) outer membrane (11–13). In contrast, the C-terminal region is responsible for making the inter-molecular contacts necessary to form the physiological S-layer lattice (10). A recent 2.7 Å resolution crystal structure revealed that the C-terminus, RsaA_249-1026_, consists of a series of beta-strands arranged in an L-shape and connected by loops interspersed with calcium-binding Repeat-in-Toxin (RTX) motifs (Figure 1A,B) (10, 14). Docking this crystal structure into a 7.4 Å resolution electron cryo-tomographic reconstruction of the repeating hexameric unit of the native S-layer lattice confirmed the identity of the side chains coordinating the physiological S-layer crystal contacts (Figure 1A and S1) (10). This S-layer reconstruction also revealed the location and overall topology of the N-terminal domain, which forms an anchoring ring beneath and around the center of each hexameric repeat made by the C-terminal crystallization domain (Figure 1A) (10, 11).

**Figure 1:**
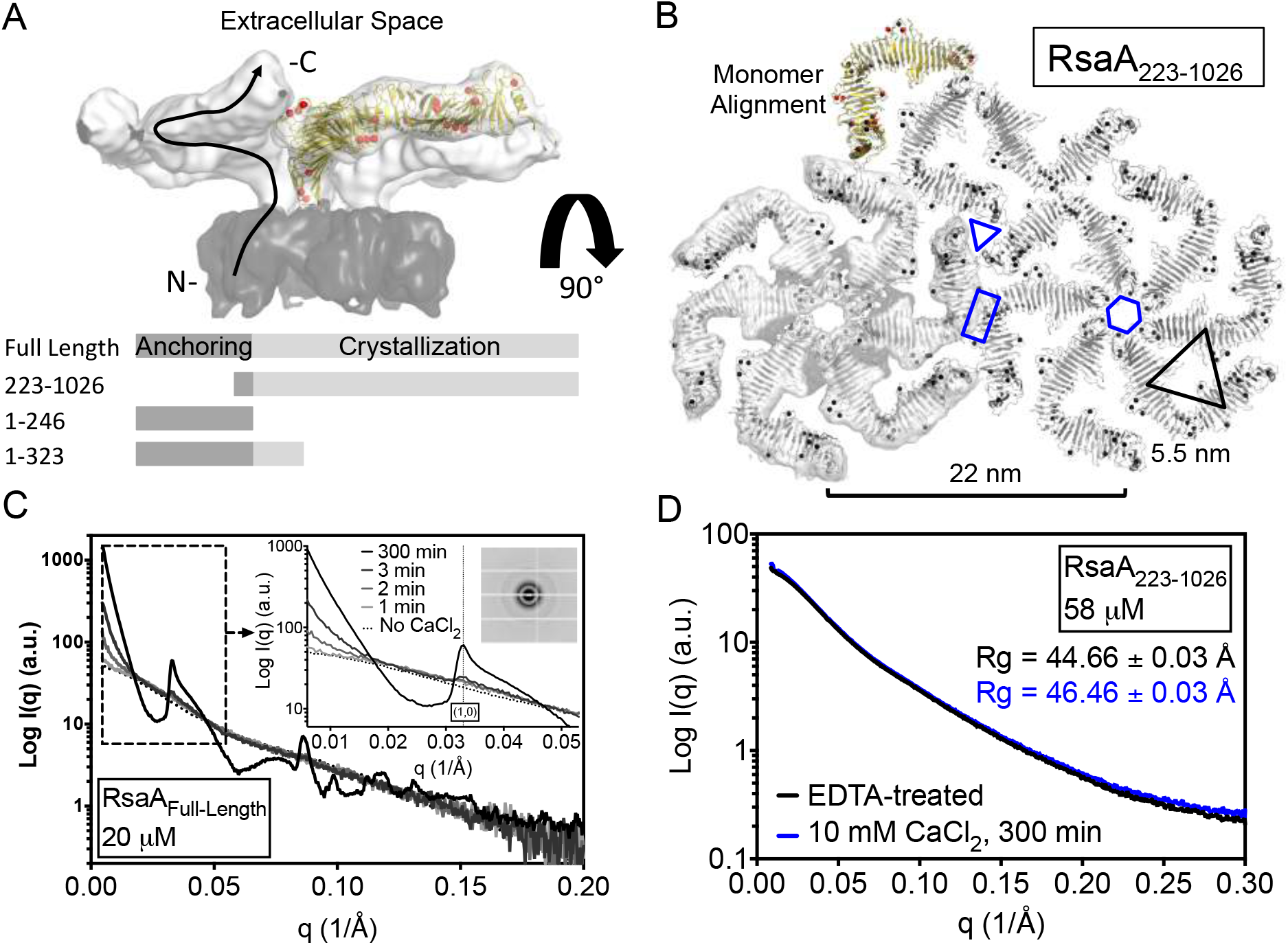
The RsaA N-terminus enhances the rate of crystallization. A) The published cryo-tomographic reconstruction of the RsaA hexameric repeat is shown with the crystallization domain surface in light grey and the anchoring domain surface in dark grey (EMD-3604). The previously determined crystal structure of the crystallization domain is docked into one arm of the hexamer with protein ribbon in gold and calcium ions as red spheres (PDB-5N97). Below, a graphical representation of the four RsaA truncation constructs used in this study is given. B) The RsaA_223-1026_ crystal structure is shown (grey ribbons with black spheres for strontium ions) with symmetry mates forming the physiological S-layer lattice. A hexamer is shown aligned with the cryo-tomographic reconstruction from (A) (grey surfaces) and an alignment of monomers from both crystal structures is shown at the top. Conserved dimeric, trimeric, and hexameric interfaces are noted by a blue rectangle, triangle, and hexagon, respectively. The 22 nm characteristic spacing between hexameric repeats is noted at the bottom and a black 5.5 nm equilateral triangle notes a pseudo-symmetric center. C) Time-resolved SAXS of 20 μM RsaAFuii-Length shows the appearance of powder diffraction when calcium is added. Bragg signal due to powder diffraction (detector image inset) appears within 2 minutes after calcium addition (plot inset marked by dashed box). D) Time-resolved SAXS of 58 μM RsaA_223-1026_ shows no evidence of self-association after 5 hours of incubation with 10 mM CaCl_2_.

Using super-resolution microscopy and single-molecule tracking, we have recently shown that during native S-layer assembly, RsaA monomers diffuse while non-covalently attached to the cell surface until incorporated into a growing crystal lattice (9). When no previous S-layer lattice exists, RsaA nucleates 2D crystals at remarkably low concentrations to form stable crystalline rafts with as few as ~50 molecules (9). Advances in imaging modalities such as atomic force microscopy (AFM) and Cryo-EM have allowed for the direct observation of macromolecular nucleation and crystallization for a select few systems at the nanometer scale (15–18). These studies have revealed canonical and non-canonical assembly pathways, including other bacterial S-layer proteins, which can exhibit structurally distinct kinetic intermediates (15, 17, 19–21). Therefore, we asked how the kinetic pathway of RsaA self-assembly might enable efficient S-layer assembly observed *in vivo*. Using rationally designed RsaA truncations (Figure 1A) and a combination of X-ray crystallography, SAXS, time-resolved Cryo-EM, stability assays, and CD spectroscopy, we identify *in vitro* structural dynamics that enable rapid 2D crystallization of the *C. crescentus* SLP, RsaA.

## Results

### The RsaA N-terminus Enhances the Rate of Crystallization

Recent attempts to determine the high-resolution structure of RsaA_Full-Length_ resulted in a 2.7 Å resolution crystal structure of the C-terminal domain (RsaA_249-1026_) (10). This 3D crystal is composed of stacked sheets of hexameric 22 nm repeats characteristic of the 2D physiological S-layer crystal lattice (10). To determine whether the unknown structure of the N-terminus contributed to the formation of stacked sheets of the S-layer lattice, RsaA_223-1026_, was cloned, purified, crystallized by the hanging drop method, and diffraction data collected to 2.1 Å resolution (22). RsaA_223-1026_ was crystallized with strontium as a calcium analog and the previously reported structure (10) was used as a search model for the molecular replacement method (Table S1). The RsaA_223-1026_ crystal structure consists of stacked hexameric sheets with substantially similar calcium-binding sites and crystal contacts within and between subunits as previously determined (Figure 1B and S1) (10). This crystal packing indicates that the crystallization domain can stably assemble into stacked sheets of the S-layer lattice without the N-terminal domain.

To compare the kinetics by which RsaAFull-Length and RsaA_223-1026_ self-assemble in the presence of CaCl_2_, we performed time-resolved SAXS on both protein forms. Previous work showed that purified RsaA_Full-Length_ crystallizes into sheets upon the addition of 10 mM CaCl_2_ (7, 8). RsaA_Full-Length_ at 20 μM exhibited Bragg peaks consistent with a 2D protein crystal lattice within two minutes of calcium addition (Figure 1C) (7). By 5 hrs, a clear powder diffraction pattern emerged, consistent with a lattice of hexameric 22 nm repeats (Figure 1C). Time-resolved SAXS of 62 and 100 μM RsaA_Full-Length_ using a stopped-flow mixer to introduce calcium quickly (within 50 ms) showed evidence of rapid assembly. At these concentrations, Bragg peaks appeared within seconds of calcium addition (Figure S2). In stark contrast, 58 μM RsaA_223-1026_ remained soluble and monomeric 5 h after calcium addition (Figure 1D). Therefore, the RsaA N-terminus enhances the rate of *in vitro* crystallization by at least three orders of magnitude.

### A Calcium-triggered Conformational Change Precedes Nucleation

Calcium-triggered self-assembly of RsaA was examined by subjecting multiple concentrations of RsaA_Full-Length_ to SAXS analysis precisely one minute after calcium addition (Figure 2A-C). Crystallization in suspension can be monitored by quantifying scattering signal in the scattering angles corresponding to the (1,0) diffraction peak of the RsaA crystal lattice (0.032<q<0.048 Å^-1^). RsaA crystallization as measured one minute after calcium addition is concentration-dependent, with initial signal appearing at 25 μM and steady-state crystal growth observed for concentrations above 35 μM (Figure 2A). To determine whether any conformational changes occur within the protein after calcium addition, but before the appearance of crystals, SAXS data from monomeric RsaA were compared to data collected one minute after calcium addition for RsaA at and below 25 μM (Figure 2B). A change in scattering intensity was observed at low scattering angles prior to the appearance of Bragg diffraction signal (Figure 2B). This signal corresponds to a decrease in the calculated radius of gyration (Rg) from ~55 Å for calcium-free monomeric protein to ~51 Å for pre-crystallized (calcium added) RsaA_Full-Length_ (Figure 2C and S3). Therefore, a calcium-triggered conformational change occurs after calcium addition, but before self-association.

**Figure 2:**
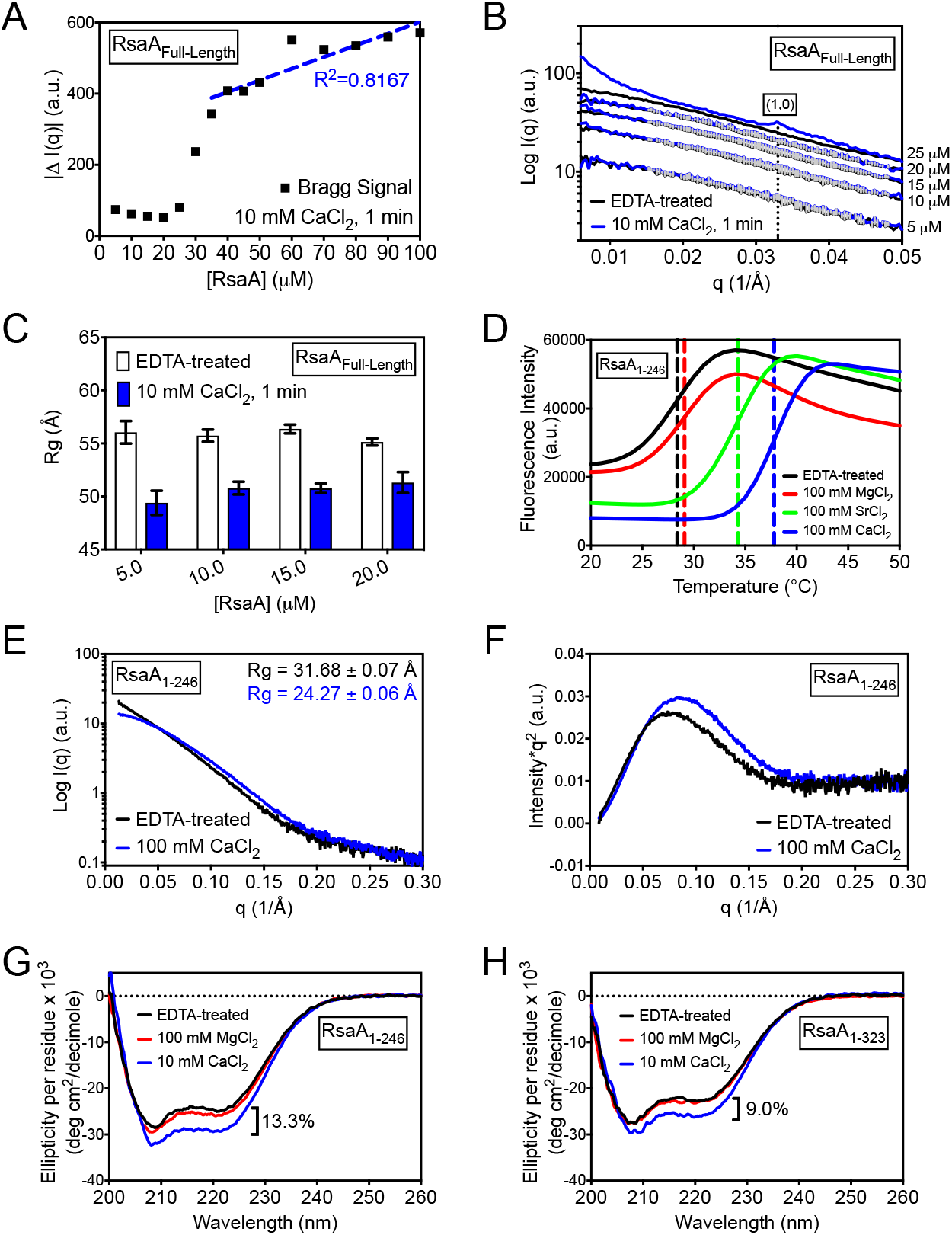
A conformational change within the N-terminus precedes nucleation. A) Bragg signal from SAXS profiles of a variety of concentrations of RsaA_Full-Length_ one minute after 10 mM CaCl_2_ addition reveals steady-state crystallization at concentrations above 35 μM. Bragg signal was measured for the (1,0) diffraction peak located at 0.032<q<0.048 Å^-1^ B) SAXS profiles of EDTA-treated (black) or pre-nucleation (10 mM CaCl_2_ for 1 min, blue) RsaA_Full-Length_ samples reveal a calcium-induced change in scattering profile. C) Differences in Rg of EDTA-treated RsaA_Full-Length_ (black) or samples 1 min after adding 10 mM CaCl_2_ (blue) show a decrease in Rg by ~4 Å before crystallization. Error bars are SEM. D) Differential scanning fluorimetry assays of RsaA_1-246_ treated with EDTA (black), 100 mM MgCl_2_ (red), 100 mM SrCl_2_ (green), or 100 mM CaCl_2_ (blue) shows calcium-specific stabilization with melting temperatures denoted by vertical dashed lines. E, F) SAXS profiles and Kratky plots of EDTA-treated (black) or 100 mM CaCl_2_-added (blue) RsaA_1-246_ shows a change in scattering profile corresponding to a decrease in Rg by 7.4 Å. G, H) Circular dichroism spectra of RsaA_1-246_ and RsaA_1-323_ are shown treated with EDTA (black), 100 mM MgCl_2_ (red), or 10 mM CaCl_2_ (blue). A decrease in the relative signal change induced by calcium indicates that the observed folding change occurs entirely within RsaA_1-246_.

### The N-terminus Increases Secondary Structure When Exposed to Calcium

SAXS profiles of RsaA_223-1026_ with and without 10 mM CaCl_2_ show no significant differences in scattering profile despite many known calcium-binding sites within this region of the protein (Figure 1B,D and S4). The N-terminal domain may then be responsible for the calcium-dependent conformational change observed in Figure 2B. Although RsaA_1-246_ lacks putative calcium binding motifs, calcium-specific stabilization was observed by differential scanning fluorimetry (Figure 2D and S5). To probe structural changes due to calcium binding, we collected SAXS profiles of RsaA_1-246_ with and without 100 mM CaCl_2_ (Figure 2E). RsaA_1-246_ displayed a significant change in scattering profile corresponding to a decrease in Rg of 7.4 A and an increase in globularity (Figure 2E, F, and S4).

Given the calcium-specific conformational change observed for RsaA_1-246_, we utilized circular dichroism (CD) spectroscopy to compare the secondary structure of the N-terminus in the presence of 5 mM EDTA, 10 mM CaCl_2_, or 100 mM MgCl_2_. A calcium-specific response was evident, corresponding to a 13.3% increase in secondary structure (Figure 2G). To determine whether this effect was confined to the N-terminus, we created a hybrid construct including all of the N-terminus and the first 7 beta-strands (77 amino acids) of the C-terminus, RsaA_1-323_ (Figure 1A). CD spectra of this construct revealed a calcium-specific 9.0% increase in secondary structure (Figure 2H), corresponding to a 32% decrease in relative signal with a 27% increase in relative mass between the two constructs. Therefore, the C-terminal 77 amino acids in RsaA_1-323_ do not contribute to the observed calcium-induced change in secondary structure, which occurs entirely within the N-terminal domain, RsaA_1-246_.

### A Short-lived Intermediate Crystal Lattice is Observed During Nucleation

To observe the steps leading to the appearance of the RsaA crystal lattice, we imaged RsaA nucleation using time-resolved Cryo-EM. Samples of 20 μM RsaA_Full-Length_ were frozen on an EM grid 30 s, 60 s, 90 s, and 120 s after adding 10 mM CaCl_2_. Multiple structural states of RsaA were observed (Figure 3). The first structure, termed the intermediate lattice, appears within 30 s after calcium addition (Figure S6) and dominates the viewing area at the 60 s time point (Figure 3A). The intermediate lattice exhibits a characteristic spacing of 5.5 nm, which may reflect pseudo-symmetric trigonal spacing formed between neighboring monomers within each repeating hexamer (Figure 1B and 3A, insets). At 90 s, the sample becomes heterogeneous, consisting of the previously observed intermediate lattice as well as the mature lattice, which exhibits the expected 22 nm spacing in Fourier space (Figure S6). At 120 s, single and stacked layers of mature lattices are predominantly observed (Figure 3B and S7). Quantification of the appearance of the intermediate and mature lattices throughout the time course reveals the rapid appearance and disappearance of the intermediate lattice within 120 s (Figure 3C). Therefore, time-resolved Cryo-EM imaging revealed a short-lived crystalline intermediate during RsaA nucleation.

**Figure 3:**
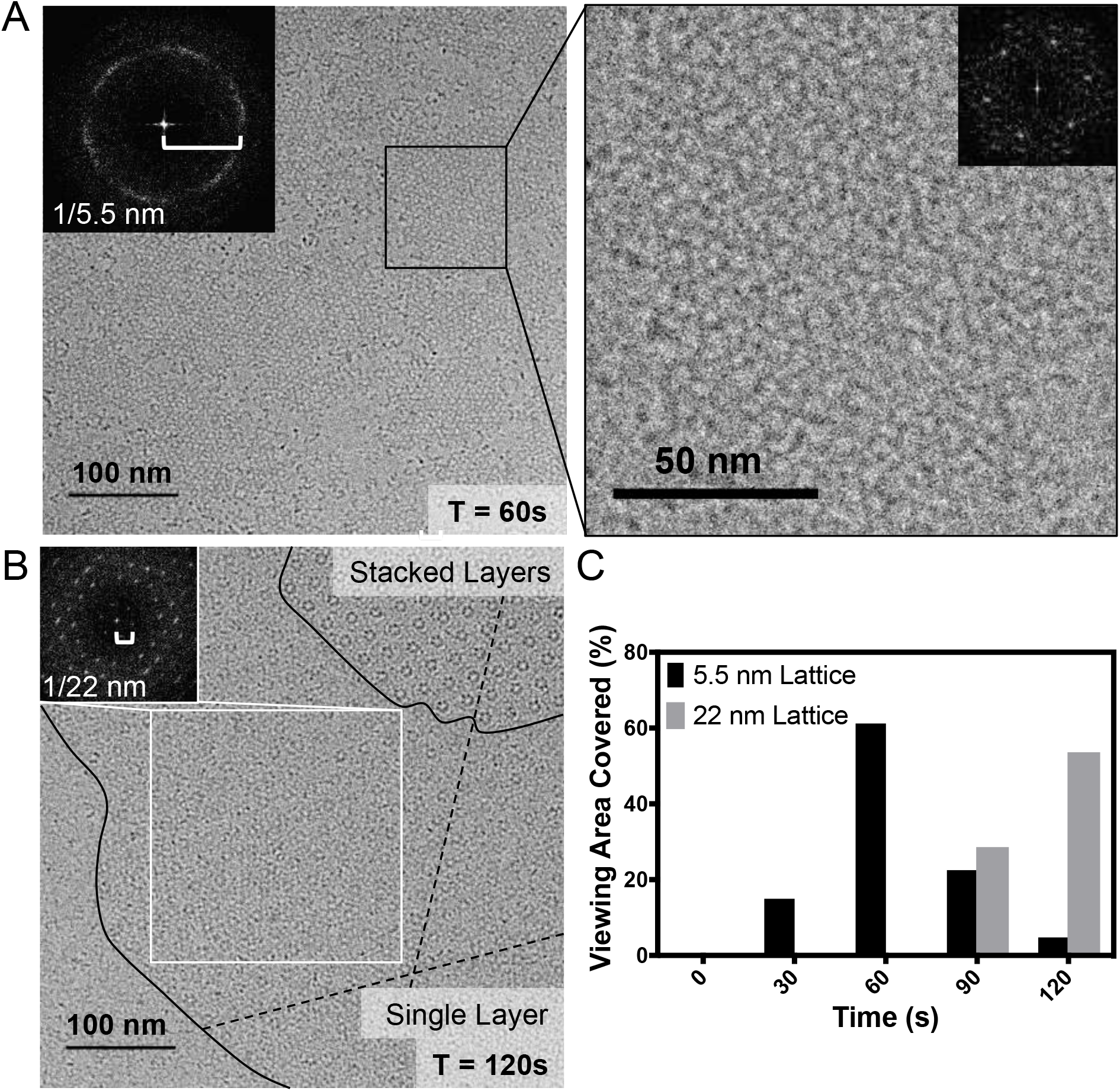
RsaA exhibits a short-lived intermediate crystal lattice during nucleation. A) A Cryo-TEM image of 20 μM RsaA_Full-Length_ 60 s after 10 mM CaCl_2_ addition shows crystal lattices with a characteristic spacing of 5.5 nm (insets are FFTs). B) A Cryo-TEM image of 20 μM RsaA_Full-Length_ 120 s after calcium addition shows a crystal lattice (noted by dashed lines) with single-layer and stacked-layer sections (boundaries noted by black lines) exhibiting a characteristic spacing of 22 nm (inset is FFT of white box). C) Quantitating the viewing area covered by 5.5 nm and 22 nm crystal lattices for at least 2.9 μm^2^ of total area at each time point shows appearance and disappearance of the 5.5 nm lattice.

### The Intermediate Crystal Lattice is Partially Ordered

To elucidate the structural differences between the intermediate and mature lattices at roughly nanometer resolution, 2D averages of the repeating hexamers composing each crystalline state were calculated. These structures were then compared to the previously determined sub-tomographic average of the physiological S-layer (EMD-3604) and a truncated version corresponding to the crystallization domain alone (using PDB-5N97 as a guide) (Figure 4A-D). The intermediate state possesses the pinwheel structure characteristic of the crystallization domain alone (Figure 4A and B) and contrasts with the physiological RsaA lattice, which possesses significant density in a ring near the center of each RsaA hexamer (Figure 1A and B). Averaging hexameric repeats from single-layer mature lattices shows electron density that more closely approximates the physiological lattice (Figure 4C and D).

**Figure 4:**
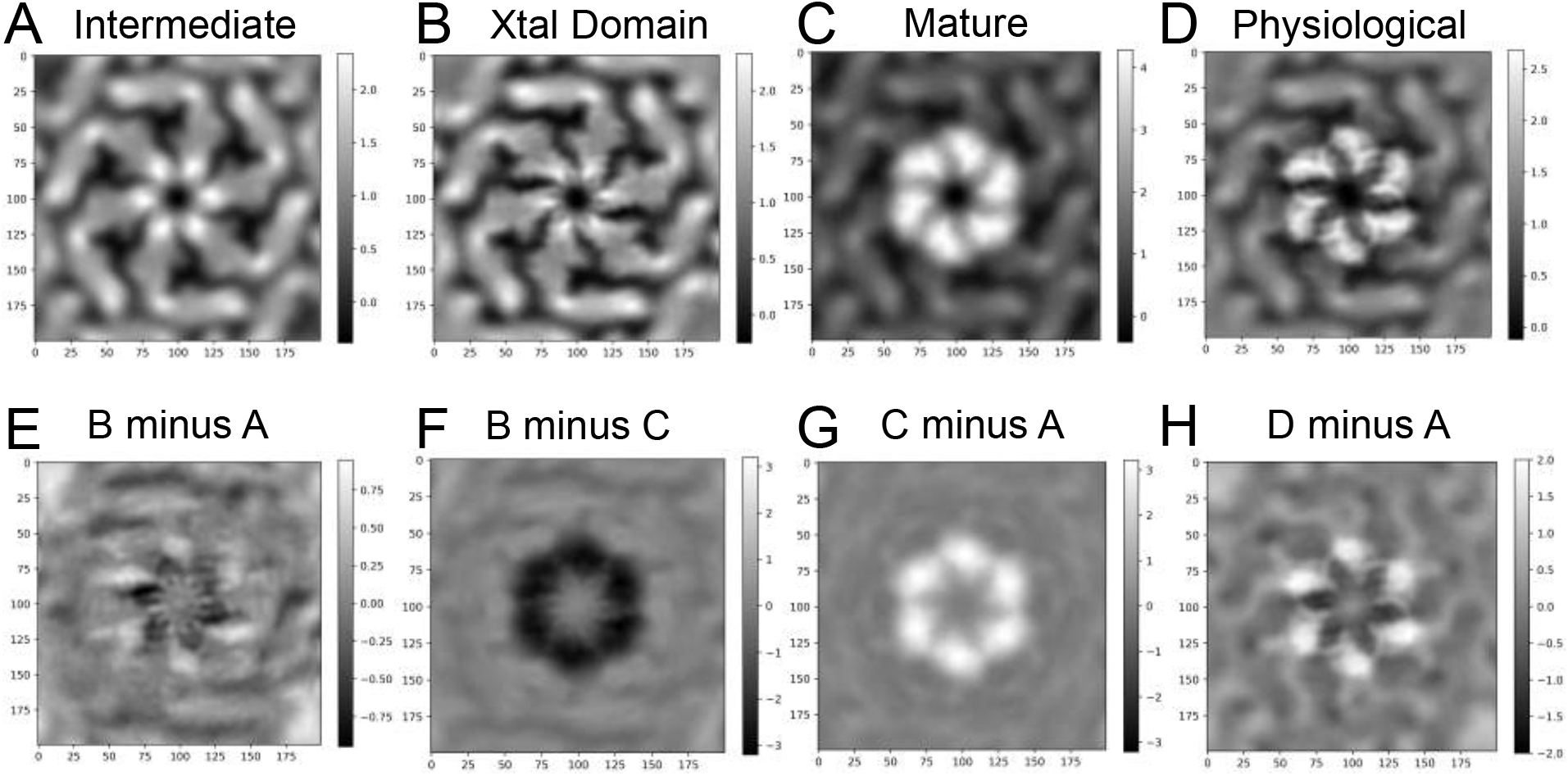
The N-terminal domain exhibits motional dynamics in the crystalline intermediate. A) A 2D average of 1044 hexameric particles from intermediate lattices as shown in Figure 3A is produced. B) The sub-tomographic average of the native crystallization domain alone is shown (EMD-3604). C) A 2D average of 1163 hexameric particles from mature single layer lattices as shown in Figure 3B is produced. D) The sub-tomographic average of the complete native hexameric repeating unit is shown (EMD-3604). E) The difference map subtracting the intermediate (A) from the crystallization domain alone (B) shows near complete signal removal. F) The difference map subtracting the time-resolved mature lattice (C) from the crystallization domain alone (B) shows negative differences around the center of the repeating hexameric unit (black). G) The difference map subtracting the time-resolved intermediate lattice (A) from the time-resolved mature lattice (C) produces positive differences around the center of the repeating hexameric unit (white). H) Subtracting the time-resolved intermediate (A) lattice from the physiological lattice (D) produces localized positive differences at the location of the N-terminal anchoring/nucleation domain (white).

To better define the structural dynamics within and between these states, difference maps were calculated between these averaged images (Figure 4E-H). Subtracting the intermediate state (Figure 4A) from the crystallization domain (Figure 4B) yields little residual signal (Figure 4E), indicating that this domain is the predominant region of the protein that is ordered during this state. Subtracting the single layer mature lattice (Figure 4C) from the crystallization domain alone (Figure 4B) creates negative differences in a ring around the center of the repeating hexameric unit (Figure 4F). Strikingly, subtracting the intermediate lattice (Figure 4A) from either the mature or physiological lattices (Figure 4C and D) produces positive electron density in a similar ring around the symmetric center of the repeating hexameric unit (Figure 4G and H). These difference maps indicate that the intermediate lattice lacks electron density at the location corresponding to the N-terminal anchoring domain. Thus, the N-terminal domain retains flexibility in the intermediate state until the subsequent mature conformation is completed.

## Discussion

Despite the vast diversity of SLPs (5, 23), the molecular interactions responsible for maintaining a physiological S-layer lattice have been elucidated by crystallography and confirmed by EM for only two proteins to date: RsaA and SbsB, the SLP from the gram-positive bacterium *Geobacillus stearothermophilus* (6, 10). Calcium-binding in SbsB was shown to directly mediate interactions between domains, thereby compacting the structure into the conformation suitable for crystallization (6). In the case of RsaA, even though calcium-binding sites are evident throughout the crystallization domain, all crystal contacts between monomers occur through amino acid interactions, not by coordinating divalent calcium ions (Figure 1B and S1) (10). This observation was supported by our SAXS analysis of RsaA_223-1026_ indicating that calcium binding does not significantly change the structure of the crystallization domain (Figure 1D). However, calcium stabilizes the secondary structure of RsaA and binds near residues directly involved in crystal contacts such as P693 and T758 at the dimeric interface and T256 at the hexameric interface (Figures S1 and S5) (7, 10).

The 2.1 Å resolution crystal structure of RsaA_223-1026_ revealed that although calcium-binding along the crystallization domain does not induce an observable conformational change, this binding behavior is sufficient to assemble stacked sheets of the physiological S-layer lattice (Figure 1B). However, this macroscopic assembly of RsaA_223-1026_ occurred over multiple days in the presence of precipitants and aided by a slow concentration process due to vapor equilibration (22). In stark contrast, upon adding 10 mM CaCl_2_ to purified RsaAFull-Length, self-assembly occurs readily as observed with small angle x-ray diffraction within two minutes at 20 μM (Figure 1C). As a direct comparison, RsaA_223-1026_ failed to self-assemble in 5 h under the same buffer conditions and a higher protein concentration (Figure 1D). Therefore, the N-terminal domain of RsaA enhances self-assembly kinetics and acts as a nucleation domain.

Monitoring protein crystal nucleation at the nano-scale by time-resolved Cryo-EM (Figure 3) revealed that after calcium binding, RsaA crystallizes into sheets via a multi-step pathway. At 20 μM, RsaA_Full-Length_ self-assembles into mature crystals within 120 s (Figure 1C and 3B) by first forming a structurally distinct short-lived crystalline intermediate at earlier time points (Figures 4 and 5). Within this intermediate state, the nucleation domain appears flexible with respect to the rigid lattice formed concurrently by the crystallization domain (Figures 4 and 5). The nucleation domain appears to reach its mature conformation by later forming a hexagonally symmetric ring around the center of each hexameric repeat made by the crystallization domain (Figures 4C and 5). This mature conformation agrees with the physiological S-layer structure, in which the nucleation domain resides beneath the crystallization domain (Figures 1A, 4D, and 5). Crystallization of SbpA, the calcium-triggered SLP from *Lysinibacillus sphaericus*, was recently observed by time-resolved AFM and exhibited a structural intermediate evidenced by discrete changes in crystal height (21, 24). Delayed assembly of specific domains, such as the N-terminal domain of RsaA (Figure 5), could be responsible for intermediate conformations observed for other SLPs like SbpA.

**Figure 5:**
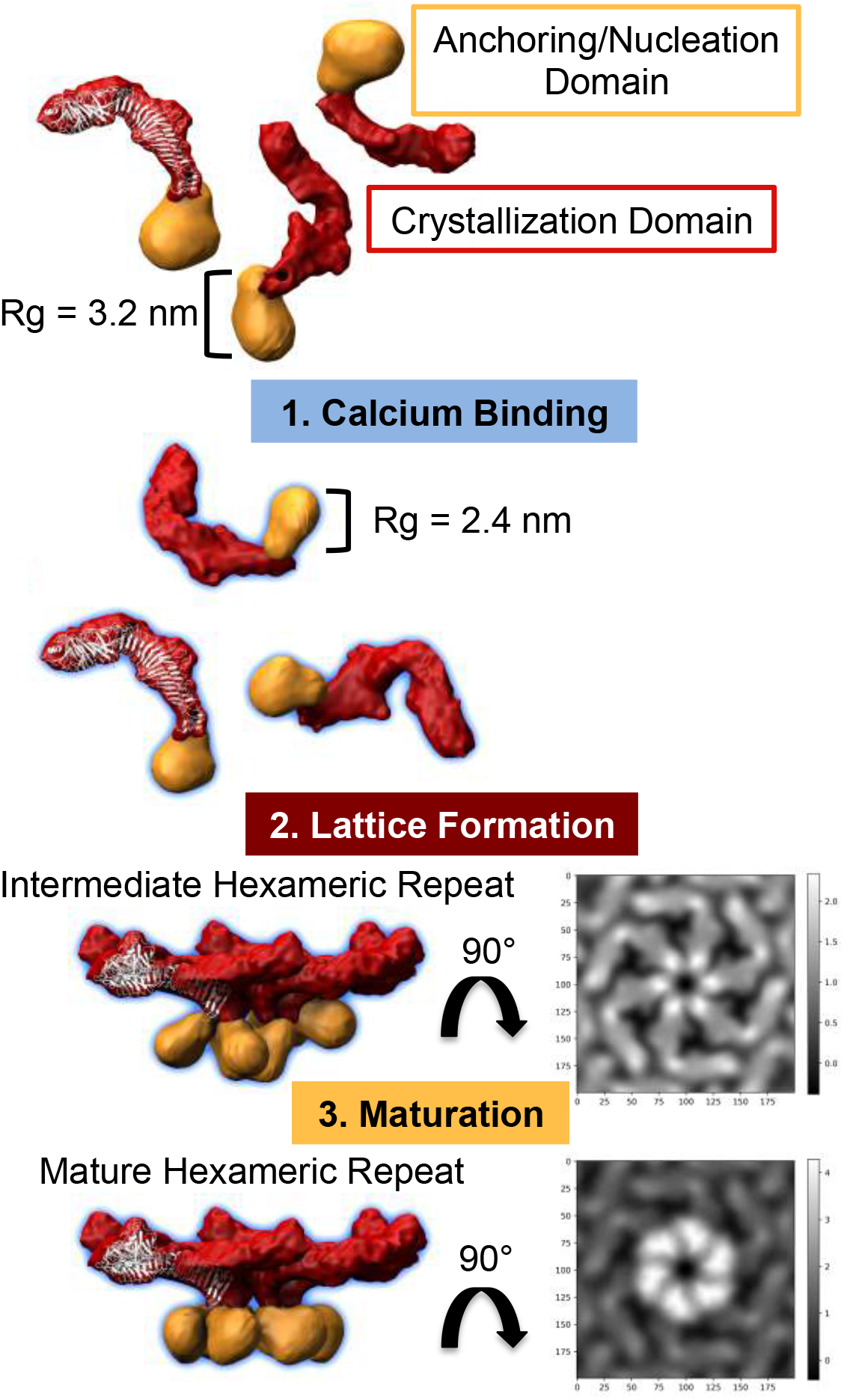
Rapid *in vitro* RsaA crystallization involves a multi-step pathway. Without calcium, the crystallization domain of RsaA (red surface, white crystal structure) is folded, but the N-terminus (orange surface) exists in a partially unfolded state. Upon calcium introduction, the N-terminus folds and compacts (Rg decreases by 7.4 Å). Then, monomers begin to assemble into a lattice using the crystallization domain. However, the N-terminus retains motion and therefore does not appear in the 2D average of the intermediate state as observed by Cryo-TEM. Later, the N-terminus locks into place as confirmed by another 2D average, forming a state resembling the physiological crystal lattice. Thus, a multi-step pathway enables rapid protein crystallization for this bacterial SLP.

The crystallization domain possesses all the residues necessary to form the long-range S-layer lattice, yet we observe a nucleation domain-dependent multi-step assembly pathway of RsaA_Full-Length_. How does the nucleation domain mediate assembly kinetics? The precise mechanism by which this nucleation domain enhances the rate of self-assembly without directly participating in most intermolecular contacts remains unclear. By isolating and examining RsaA truncations, we determined that the nucleation domain undergoes a calcium-specific conformational change before crystallization, likely mediated by specific structural changes including increasing helical content of the domain upon binding calcium (Figure 2). This observed calcium-induced conformational change might affect the formation of the hexameric interface of the crystallization domain, which is just downstream in the RsaA sequence. This change may also expose additional intermolecular interfaces to be utilized during delayed maturation of the nucleation domain. Segregating the biochemical steps in this assembly pathway would likely require high-resolution structural knowledge of the nucleation domain to guide rational mutagenesis. Alternatively, in the context of a short-lived crystalline intermediate, flexibility between domains may provide access to a larger landscape of conformational microstates, which could act as an additional entropic driving force for successful crystal nucleation. A careful study of the energetics of RsaA self-assembly may be required to fully elucidate the mechanism of nucleation rate enhancement observed herein.

RsaA self-assembles quickly and efficiently *in vivo* at extremely low concentrations, (9) suggesting an evolutionary basis for non-canonical crystallization kinetics if the same or similar assembly pathway occurs on the cellular surface. SLPs continuously crystallize on the curved and growing LPS outer membrane and the topology of the surface has been shown to affect S-layer crystal growth *in vivo* (9). Flexibility between domains might allow the crystallization domain to position itself for lattice formation while relieving torque originating from anchoring a 2D lattice on a variably curved surface. Our results further indicate that SLPs may segregate kinetic regulation of assembly using a structurally remote nucleation domain that enables fast and robust crystallization through an intermediate. This modularity raises the possibility of designing biologically inspired, self-assembling macromolecular nanomaterials with controllable nucleation kinetics.

## Methods

### Protein Purification

Purified RsaA_Full-Length_ in the absence of CaCl_2_ was previously shown to partially unfold at 28°C (7). Therefore, all RsaA samples were kept cold (<4°C) at all times unless otherwise noted. RsaA_Full-Length_ protein was purified similarly to previously reported methods (7). *C. crescentus* NA1000 cells were grown to early stationary phase at 30°C in PYE medium, shaking at 200 rpm. The culture was then pelleted by centrifugation and stored at -80°C. Approximately 1 g of cell pellet was thawed on ice, re-suspended with 10 mL of ice cold 10 mM HEPES buffer pH 7.0, and centrifuged for 4 min at 18,000 rcf. This washing step was performed three times. The pellet was then separated into 10 aliquots and each was re-suspended with 600 μL of 100 mM HEPES buffer pH 2.0. These cell suspensions were incubated on ice for 15 min and then spun for 4 min at 18,000 rcf. The supernatants were then pooled and neutralized (pH = 7) by the addition of 5 N NaOH. To remove free divalent cations, 5 mM ethylenediaminetetraacetic acid (EDTA) was added. The protein solution was then syringe filtered using a 0.22 μm PES syringe filter and 5 mL were injected onto a Highload Superdex200 16/600 size exclusion column (GE Healthcare). During size exclusion chromatography, the running buffer consisted of 50 mM Tris/HCl pH 8.0 and 150 mM NaCl. Monomeric RsaA_Full-Length_ eluted at approximately 0.52 column volumes (CV) (Figure S8). From 1 g of pelleted cells, we consistently purified at least 1 mg of monomeric RsaA_Full-Length_ protein. Purity was assessed by SDS-PAGE (Figure S8).

To purify RsaA_223-1026_, a previously reported *C. crescentus* strain was obtained from the Smit laboratory (University of British Columbia) and utilized (22). Briefly, RsaA_223-1026_ was cloned into a pUC8CVX plasmid and introduced to background strain JS1014 by electroporation. JS1014 includes genomic knockouts of *rsaA, sap* (an S-layer associated protease), and *manB* (eliminating major surface polysaccharides), resulting in the secretion of RsaA_223-1026_ into the growth medium upon transformation. RsaA_223-1026_-producing cells were grown for 72 hrs in 250 mL of PYE medium. This culture was performed in a 2.5 L wide-bottom flask without shaking, to provide aeration for cell growth but limit agitation, which causes macroscopic aggregation of RsaA_223-1026_. Cultures were collected and centrifuged at 9000 rcf for 20 min to remove intact cells. The supernatant was collected, filtered using a 0.22 μm PES syringe filter, and concentrated using a 30 kDa MWCO centrifugal concentrator (Sartorius). In the same concentrator, the protein solution was buffer exchanged into 50 mM Tris/HCl pH 8.0 and 150 mM NaCl. To chelate divalent ions, 5 mM EDTA was added before injecting the solution onto a Highload Superdex200 16/600 size exclusion column with the same running buffer as for RsaA_Full-Length_. RsaA_223-1026_ eluted at 0.57 CV and purity was assessed by SDS-PAGE (Figure S8).

To purify RsaA_1-246_ and RsaA_1-323_, both sequences were cloned into pET-28a using the Nde1 and Nhe1 restriction sites, resulting in an N-terminal 6XHis-tag for affinity purification. *Escherichia coli* BL21 (DE3) cells were transformed with the plasmid encoding either RsaA_1-246_ or RsaA_1-323_. Cells were grown in LB with 30 μg/mL kanamycin, shaking at 200 rpm, until an OD_600_~0.8 was reached, at which point protein expression was induced by the addition of 0.5 mM isopropyl 1-thio-β-D-galactopyranoside (Gold Biotechnology). Temperature was reduced to 16°C for overnight expression. Cells were then pelleted and either stored at -80°C for future use or re-suspended in lysis buffer containing 50 mM Tris/HCl pH 8.0, 500 mM NaCl, 10% glycerol, and 0.2% Tween 20. Cells were lysed using a sonicator (Qsonica) and subsequently centrifuged at 31,000 rcf for 45 min. RsaA_1-246_ and RsaA_1-323_ were found to form insoluble inclusion bodies, which were isolated as the pellet from the previous spin and re-solubilized using denaturation buffer consisting of 50 mM Tris/HCl pH 8.0, 150 mM NaCl, 100 mM CaCl_2_, and 6 M guanidine/HCl. After gentle agitation for 1 hr at room temperature, the solubilized protein was centrifuged at 31,000 rcf for 45 min. The resulting supernatant was isolated and filtered using a 0.22 μm PES syringe filter. Ni-NTA resin was added to the protein solution and the resultant slurry was allowed to dialyze overnight at 4°C against 50 mM Tris/HCl pH 8.0, 150 mM NaCl, and 100 mM CaCl_2_. The next day, the Ni-NTA resin was isolated in a gravity column and washed with 50 mM Tris/HCl pH 8.0, 150 mM NaCl, and 20 mM imidazole until the flow-through was clear. Soluble, refolded protein was then eluted from the Ni-NTA resin using 10-15 mL of 50 mM Tris/HCl pH 8.0, 150 mM NaCl, and 200 mM imidazole. The resulting protein solution was concentrated to 5 mL, treated with 5 mM EDTA, and injected onto a Highload Superdex200 16/600 size exclusion column with the same running buffer as for the other constructs: 50 mM Tris/HCl pH 8.0 and 150 mM NaCl. RsaA_1-246_ and RsaA_1-323_ eluted at 0.71 CV and 0.63 CV, respectively and purity was assessed by SDS-PAGE (Figure S8).

### X-ray Crystal Structure Solution and Refinement

Diffraction data of RsaA_223-1026_ were previously collected and reported (22). A dataset collected at 0.9795 Å wavelength was reprocessed using Aimless (25) to 2.1 Å resolution based on a moderately conservative CC1/2 score of 0.6. CC1/2, the correlation of one random half of the observations to the other half, was used to determine the high resolution limit over the widely used R_merge_ and Mean(I)/σ(Mean(I)) due to more recent empirical evidence by Karplus and Diederichs (26) indicating that the latter protocols discard too much useful data. The crystal structure of RsaA_223-1026_ was solved by molecular replacement (PHENIX) using chain A from PDBID: 5N8P. Six molecules corresponding to residues 245-1026 were placed in the asymmetric unit and were iteratively modeled (Coot) and refined (PHENIX) against the data. Anomalous difference maps were generated from the previously reported 1.5421 Å wavelength dataset (22) and one collected at 0.7685 Å wavelength, both from the same crystal as the 0.9795Å wavelength dataset used to build the reported model. Phases from the model were used alongside ligand stereochemistry considerations to identify 124 strontium and 150 iodide ions. Refinement and modeling statistics can be found in Table S1. Map and coordinate files are available as entry 6P5T in the Worldwide Protein Data Bank.

### Small Angle X-ray Scattering/Diffraction

Snapshot and time-resolved SAXS experiments on RsaA_Full-Length_ were performed at the bio-SAXS beamline 4-2 at Stanford Synchrotron Radiation Lightsource (SSRL) (27). Data were collected using a Pilatus3 X 1M detector (Dectris AG) with a 3.5 m sample-to-detector distance and beam energy of 11 keV (wavelength λ=1.13 Å). SAXS data were measured in the range of 0.0045 Å^-1^ ≤ q ≤ 0.30 Å^-1^ (q = 4πsin(θ)/ λ with 2θ being the scattering angle). Snapshot SAXS data taken 1 min after adding 10 mM CaCl_2_ were collected in a series of ten or twelve 1 s exposures. Fast (seconds to sub-second) time-resolved SAXS experiments were performed using a customized SFM-400 four syringe stopped-flow mixer (Bio-Logic Science Instruments) to facilitate fast mixing. Time-resolved data sets were collected with increased x-ray beam flux using a double multilayer monochromator with a 2% energy bandpass to enable sub-second exposure times at reasonable signal-to-noise levels.

For RsaA_1-246_ and RsaA_223-1026_, SAXS data were collected at the SIBYLS beamline 12.3.1 at the Advanced Light Source (ALS) (28). In this case, a 2 m sample-to-detector distance and beam energy of 11 keV (λ=1.13 Å) allowed for a data range of 0.0086 Å^-1^ ≤ q ≤ 0.40 Å^-1^. SAXS data were collected in a series of 32 0.3 s exposures. At both light sources, the q scale was calibrated with silver behenate powder and RsaA solution aliquots were injected directly into a temperature-controlled (10°C) flow cell. All images were then reduced as previously described (28) or using the program SasTool (http://ssrl.slac.stanford.edu/~saxs/analysis/sastool.htm). Profiles of each frame were analyzed for possible effects of radiation damage and averaged automatically using SasTool or manually using Primus (29). Averaged buffer curves were subtracted from all averaged protein curves and Guinier analysis was performed using Primus (29).

### Differential Scanning Fluorimetry

Stability measurements of purified RsaA protein were performed using the Thermofluor assay, which consisted of 45 μL of protein solution between 0.2 mg/mL and 1 mg/mL mixed with 5 μL of 10X ion solution (or water) as well as 0.5 μL of 10,000X SYPRO Orange Protein Gel Stain (excitation/emission wavelength, λ = 490/590 nm) (Thermo Fisher Scientific). Temperature was increased at a rate of 1°C per minute from 4°C to 100°C and fluorescence was measured every minute with a qPCR thermocycler in FRET mode (Bio-Rad). Aggregation temperature was determined by locating the global minimum of the second derivative of the raw data. Binding data were fit to a single-site binding model using Prism (GraphPad).

### Circular Dichroism Spectroscopy

Circular Dichroism (CD) measurements were performed using a J-815 Circular Dichroism Spectrometer (Jasco). Far-UV spectra (200-260 nm) were recorded in a 1 mm path-length quartz cell with an exposure time of 1 sec/nm. The sample cell was maintained at 10°C and three scans were collected and averaged for each sample. RsaA_1-246_ and RsaA_1-323_ were brought to a final concentration of 1-3 μM and a uniform volume of CaCl_2_, MgCl_2_, or buffer was added to each sample before acquiring spectra. A buffer spectrum with the appropriate added ion was subtracted from each sample spectrum before plotting.

### Electron Microscopy

Time-resolved Cryo-EM was performed using a 200 keV TF20 electron microscope (Thermo Fisher Scientific) equipped with a K2 direct electron detector (Gatan). To begin crystallization, 10 mM CaCl_2_ was added to a solution of 20 μM RsaA_Full-Length_. At 95% humidity, 3 μL of crystallizing sample was applied to a 200 mesh, lacey carbon, copper EM grid (EMS), blotted for 2 seconds, and plunge-frozen into liquid ethane using a Leica EM GP (Leica Microsystems). Sample application and subsequent plunge freezing were carefully timed such that the EM grid entered liquid ethane 30, 60, 90, or 120 s after initial calcium addition. Batches of images totaling 3.76, 2.94, 4.71, and 4.68 μm^2^ of imaged area respectively were acquired from each time point at 29,000X magnification and 3 μm defocus.

Motion correction was performed using Digital Micrograph software (Gatan). Subsequent CTF correction and 2D averaging was performed using Relion 2.1.0 utilizing a GPU to accelerate processing. For 2D averaging, we first manually picked ~100 visible particles on an arbitrarily selected micrograph with particle diameter set to 220 Å, from which we generated an initial 2D average map. We then used the 2D average as a template to search for all candidate particles across all the available micrographs (mature: 18 micrographs; intermediate: 43 micrographs) by using the “Auto-picking” function in Relion with picking threshold 0.9 and minimum inter-particle distance 210 Å. We performed manual screening to remove stacked-layer particles and other false positives. Particles that passed the screening were then extracted from the micrographs and classified by the “2D classification” function in Relion. We used default parameters except the mask diameter was set to 450 Å and with an in-plane angular sampling of 5.625°. The final 2D average of the intermediate lattice included 1044 particles and has an estimated resolution of 11.5 Å (Figure 4A). The final 2D average of the single-layer mature lattice included 1163 particles and has an estimated resolution of 6.8 Å (Figure 4C). The 2D average of stacked-layers from the time resolved data set included 55 particles with an estimated resolution of 20.5 Å while the average from stacked-layer crystals grown overnight included 388 particles with an estimated resolution of 7.3 Å (Figure S7). To create difference maps, contrast was normalized using the intensity of the C-terminal arms of the RsaA hexameric repeat.

## Data Availability Statement

Supplementary information figures (8) are provided with this manuscript, which include representative images from all data sets and time points. Full data sets and materials are available from the corresponding author S.W. upon reasonable request.

## Acknowledgements

This work was supported in part by the National Institute of General Medical Sciences: Grant No. R35-GM118071 to L.S., and in part by the US Department of Energy, Laboratory Directed Research and Development under contract No. DE-AC02-76SF00515 and Biology and Environmental Research to S.W. L.S. is a Chan Zuckerberg Biohub Investigator. J.S. and M.E.P.M. acknowledge support by the Natural Sciences and Engineering Research Council of Canada Discovery Program (Grant Nos. RGPIN 36574-11 and 04802-15). J.H. was supported in part by the National Science Foundation Graduate Research Fellowship Program (NSF-GRFP) and the US Department of Energy Office of Science Graduate Student Research Program (DOE-SCGSR). SAXS data was collected at SIBYLS beamline 12.3.1 at the Advanced Light Source (ALS). SAXS data collection at SIBYLS is funded through DOE BER Integrated Diffraction Analysis Technologies (IDAT) program and NIGMS grant P30 GM124169-01, ALS-ENABLE. Use of the Stanford Synchrotron Radiation Lightsource (SSRL), SLAC National Accelerator Laboratory, is supported by the U.S. Department of Energy, Office of Science, Office of Basic Energy Sciences under contract No. DE-AC02-76SF00515. The SSRL Structural Molecular Biology Program is supported by the DOE Office of Biological and Environmental Research, and by the National Institutes of Health, National Institute of General Medical Sciences (including grant No. P41GM103393). The authors thank Greg Stewart/SLAC for graphic design support and Dong-Hua Chen and Kathryn Burnett for technical support.

## Author Contributions

Conceptualization: J.H., J.S., and S.W. Investigation: J.H. and F.J. Formal analysis: J.H., P.L., and A.C.K.C. Resources: I.R., T.M., and T.W. Writing—original draft: J.H. Writing—review and editing: All authors. Supervision and funding acquisition: L.S., J.S., M.E.P.M., and S.W.

## Competing Interests

The authors declare no competing financial interests.

Correspondence and requests for materials should be addressed to S.W.

## Supplementary Information (SI)

**Table S1.**
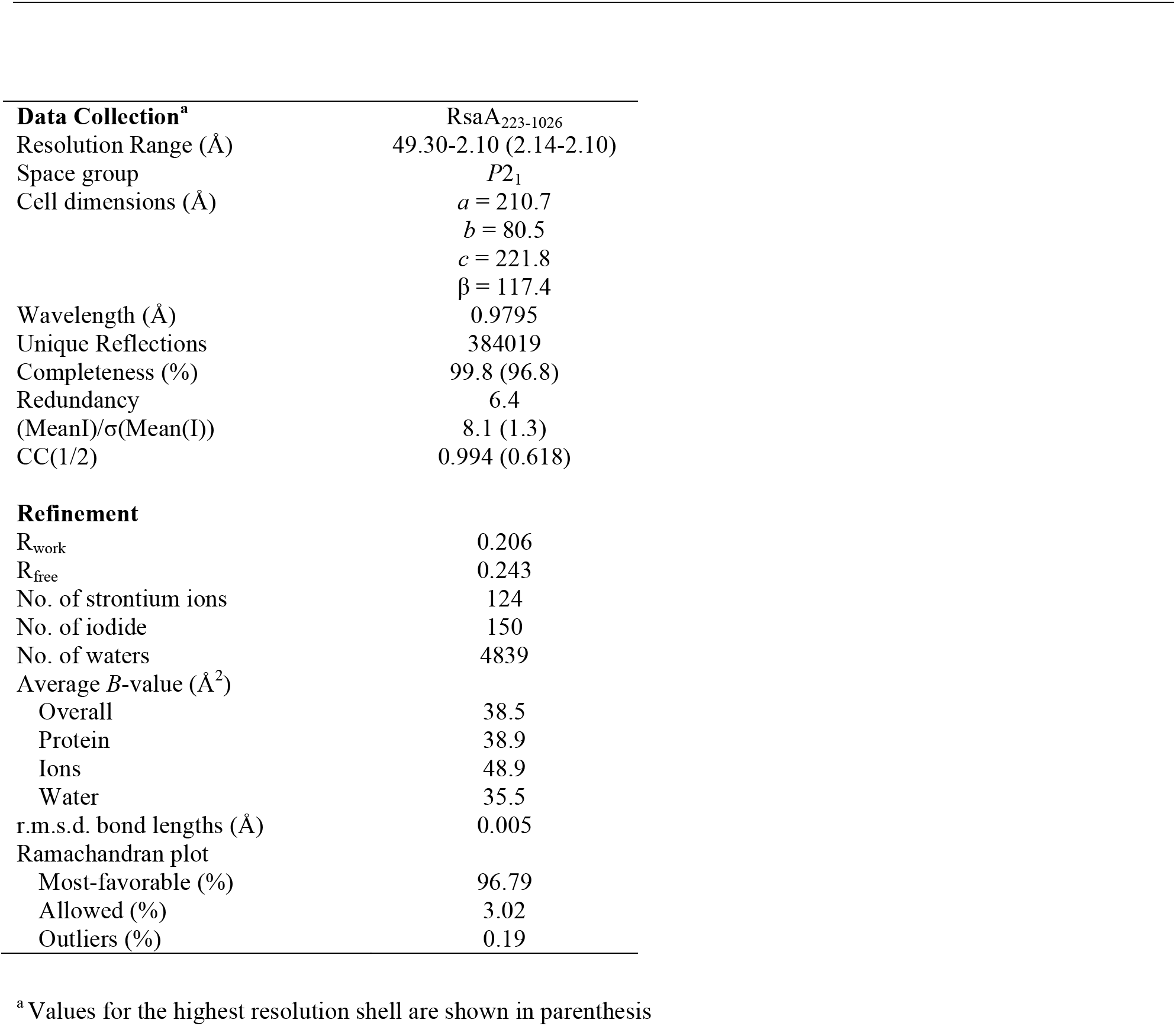
Data collection and refinement statistics.

**Figure S1:**
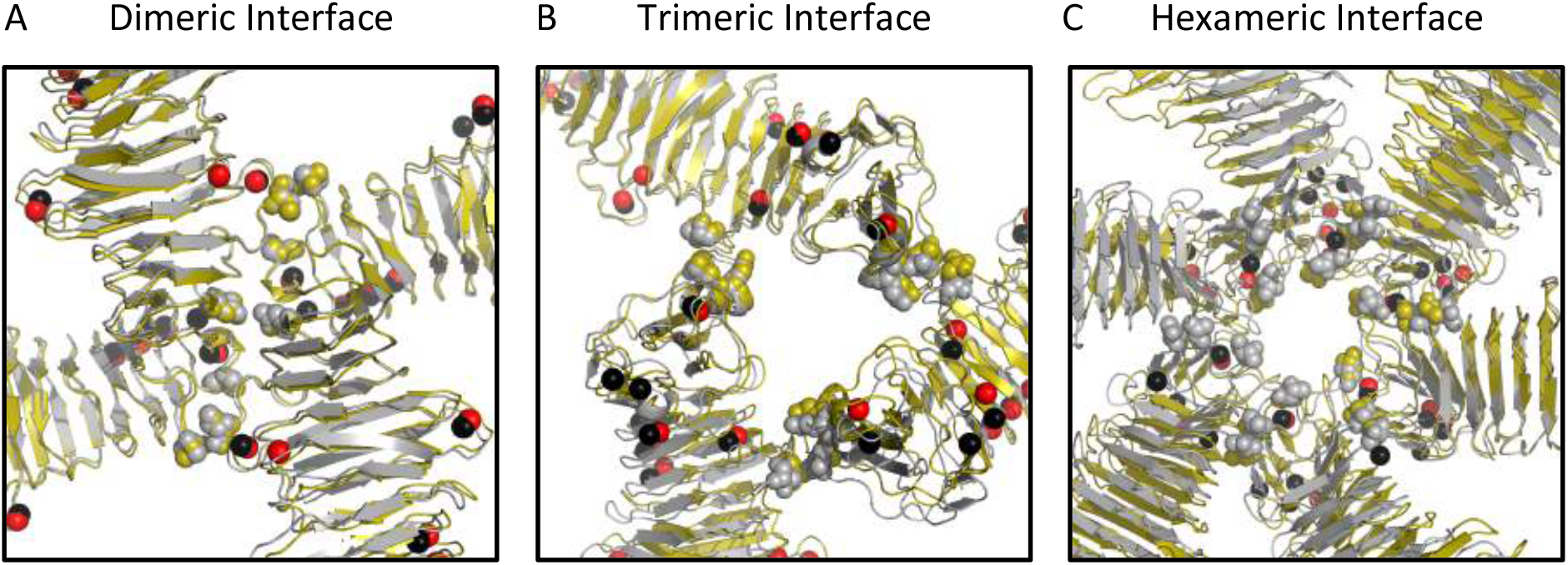
Physiological S-layer intermolecular contacts are observed in the RsaA_223-1026_ crystal structure. A, B, and C) The previously determined crystal structure (gold ribbons, calcium as red spheres, PDB-5N8P) is overlaid with the 2.1 Å resolution crystal structure of RsaA_223-1026_ (grey ribbons, strontium as black spheres). The A) dimeric, B) trimeric, and C) hexameric interfaces of both crystal structures are shown and are mediated by previously determined side chains (grey and gold spheres). Residues at the dimeric interface shown as spheres are: A667, P693, N713, and T758. Residues at the trimeric interface shown as spheres are: T854, Q960, and L952. Residues at the hexameric interface shown as spheres are: T256, V283, and T309.

**Figure S2:**
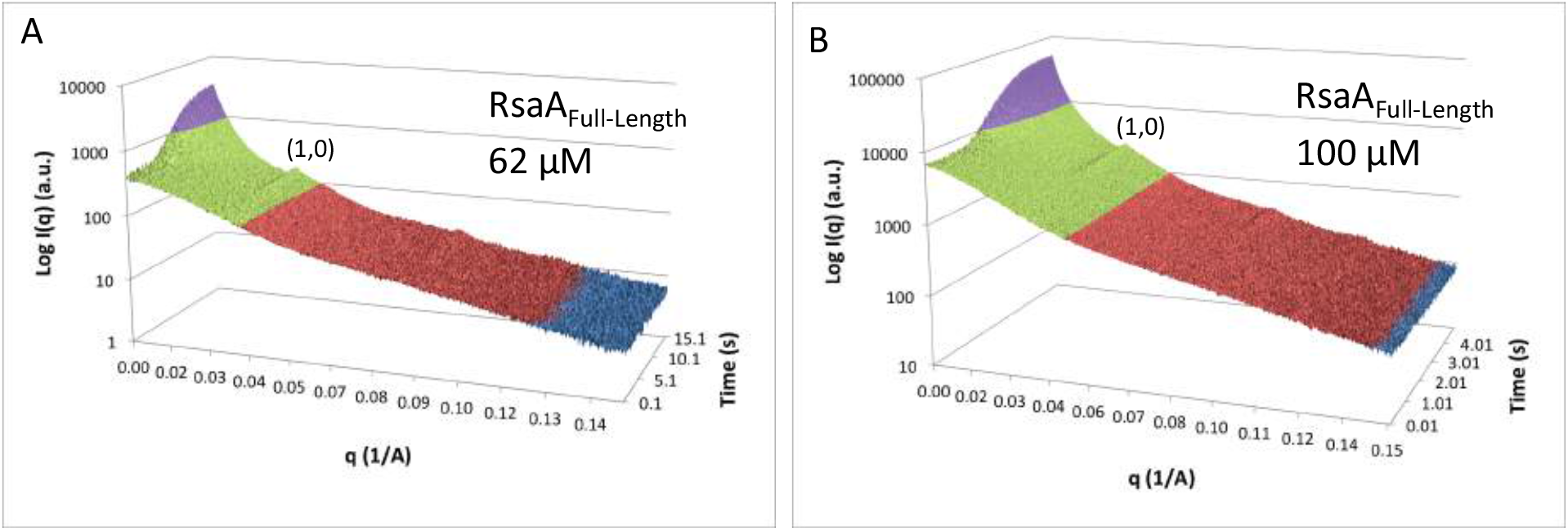
RsaA_Full-Length_ undergoes rapid calcium-triggered assembly. Time-resolved stopped-flow small-angle x-ray scattering/diffraction of A) 62 μM RsaA_Full-Length_ and B) 100 μM RsaA_Full-Length_ shows crystallization occurring within seconds at high protein concentrations. Scattering intensity is shaded from low to high (blue to red to green to purple). The (1,0) diffraction peak for the 22 nm repeat crystal lattice is noted.

**Figure S3:**
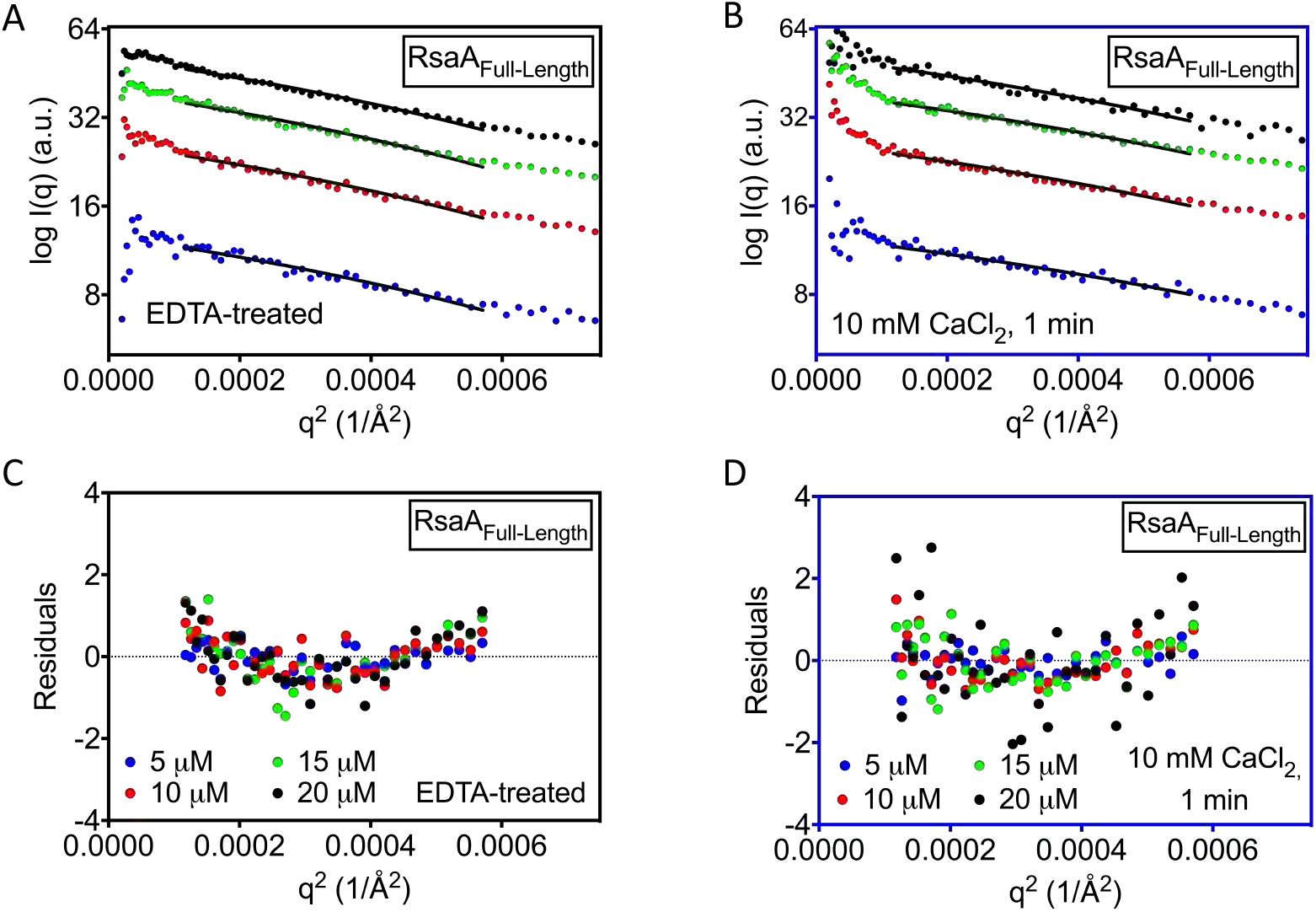
Guinier analysis of RsaA_Full-Length_ without (black frames) and with (blue frames) 10 mM CaCl_2_. A, B) Linear fits to the Guinier regime for RsaA_Full-Length_ are shown A) without calcium or B) 1 minute after adding 10 mM CaCl_2_. C, D) Residuals are plotted for the linear fits shown in A and B. In all plots, protein concentrations are 5 μM (blue), 10 μM (red), 15 μM (green), and 20 μM (black).

**Figure S4:**
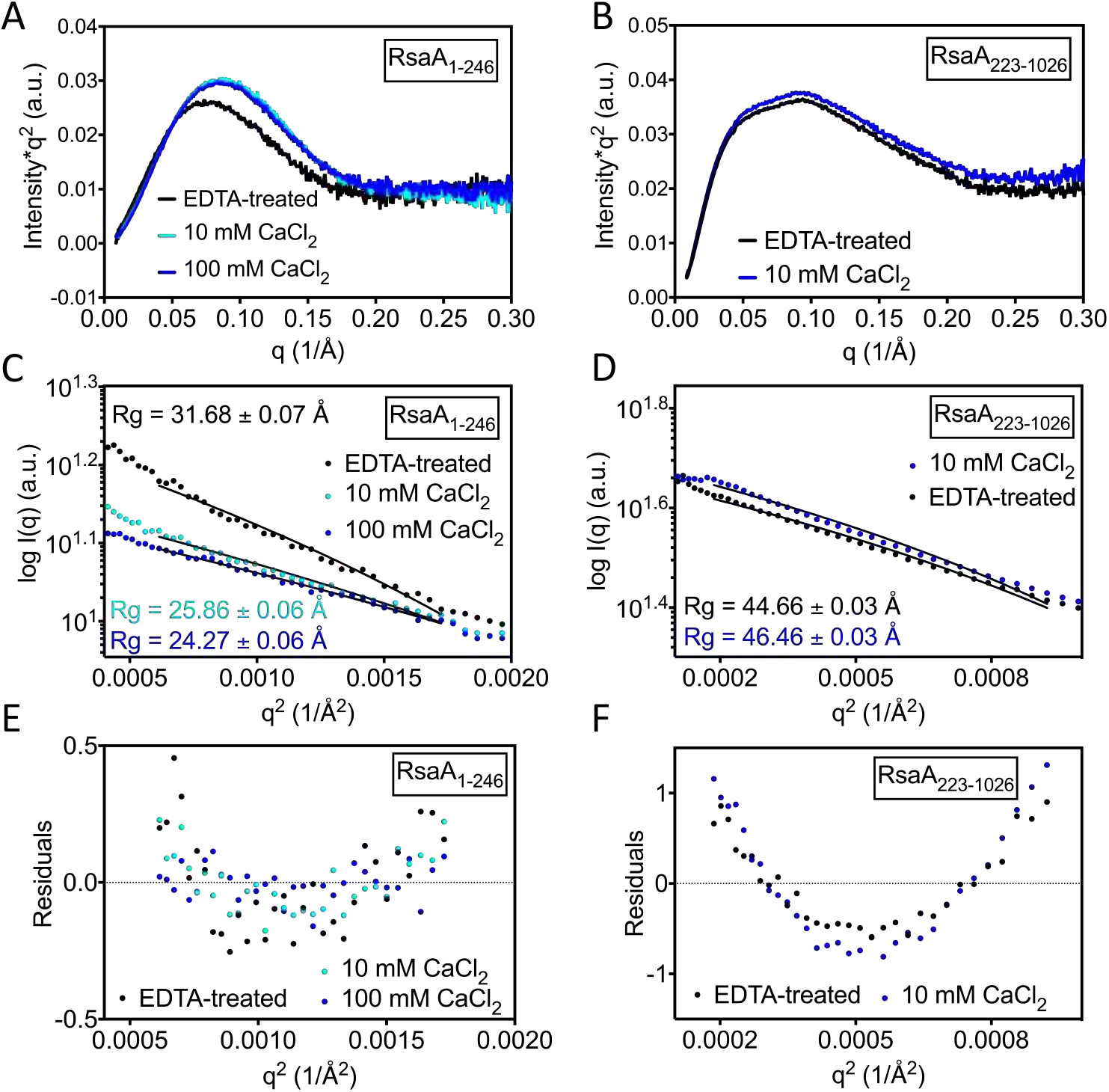
Guinier analysis of RsaA_1-246_ and RsaA_223-1026_ with and without CaCl_2_. EDTA-treated (black), 10 mM CaCl_2_ (cyan), and 100 mM CaCl_2_ samples of RsaA_1-246_ are shown in A, C, and E) while EDTA-treated (black) and 10 mM CaCl_2_ (blue) samples of RsaA_223-1026_ are shown in B, D, and F). A, B) Kratky plots of the raw SAXS data for RsaA_1-246_ and RsaA_223-1026_. C, D) Linear fits to the Guinier regime for the same samples as in A and B are shown along with the calculated Rg for each profile. E, F) Residuals are plotted for the linear fits shown in C and D.

**Figure S5:**
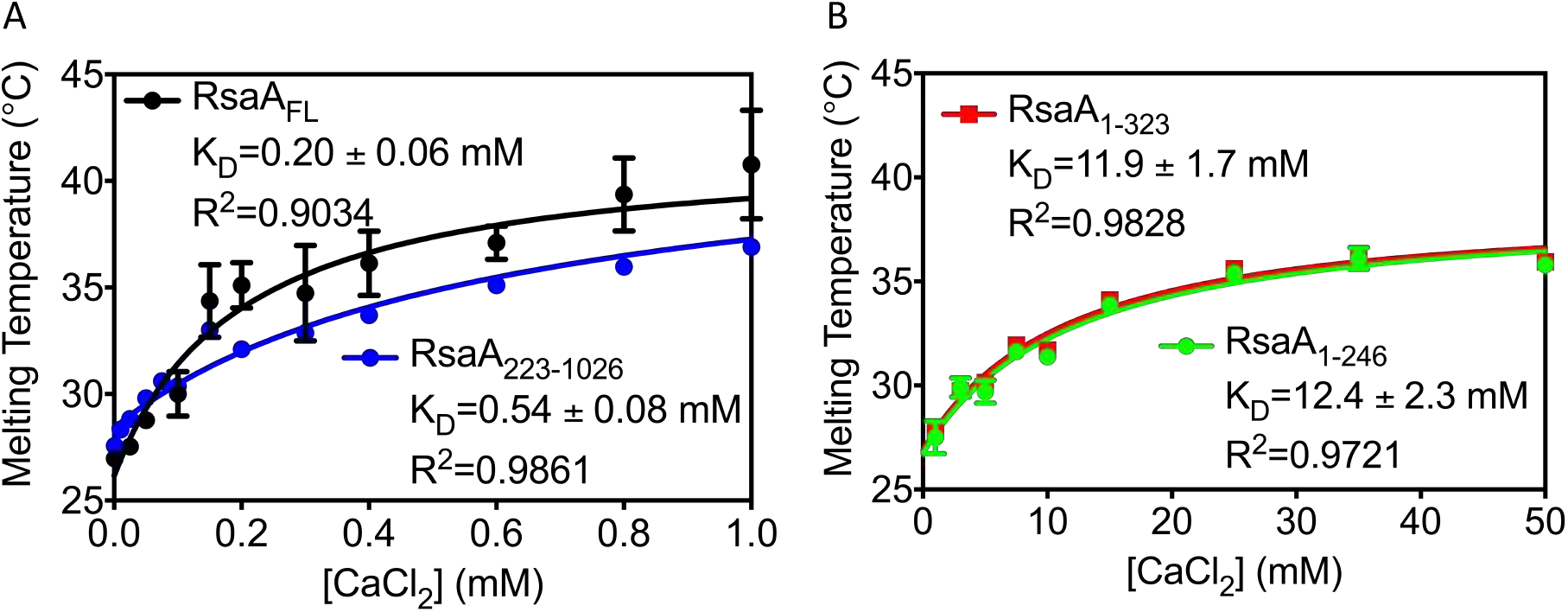
Calcium-stabilization curves for RsaA as measured by differential scanning fluorimetry. A) RsaA_Full-Length_ (black) and RsaA_223-1026_ (blue) exhibit stabilization K_D_ values less than 1 mM. B) RsaA_1-323_ (red) and RsaA_1-246_ (green) exhibit stabilization K_D_ values around 12 mM.

**Figure S6:**
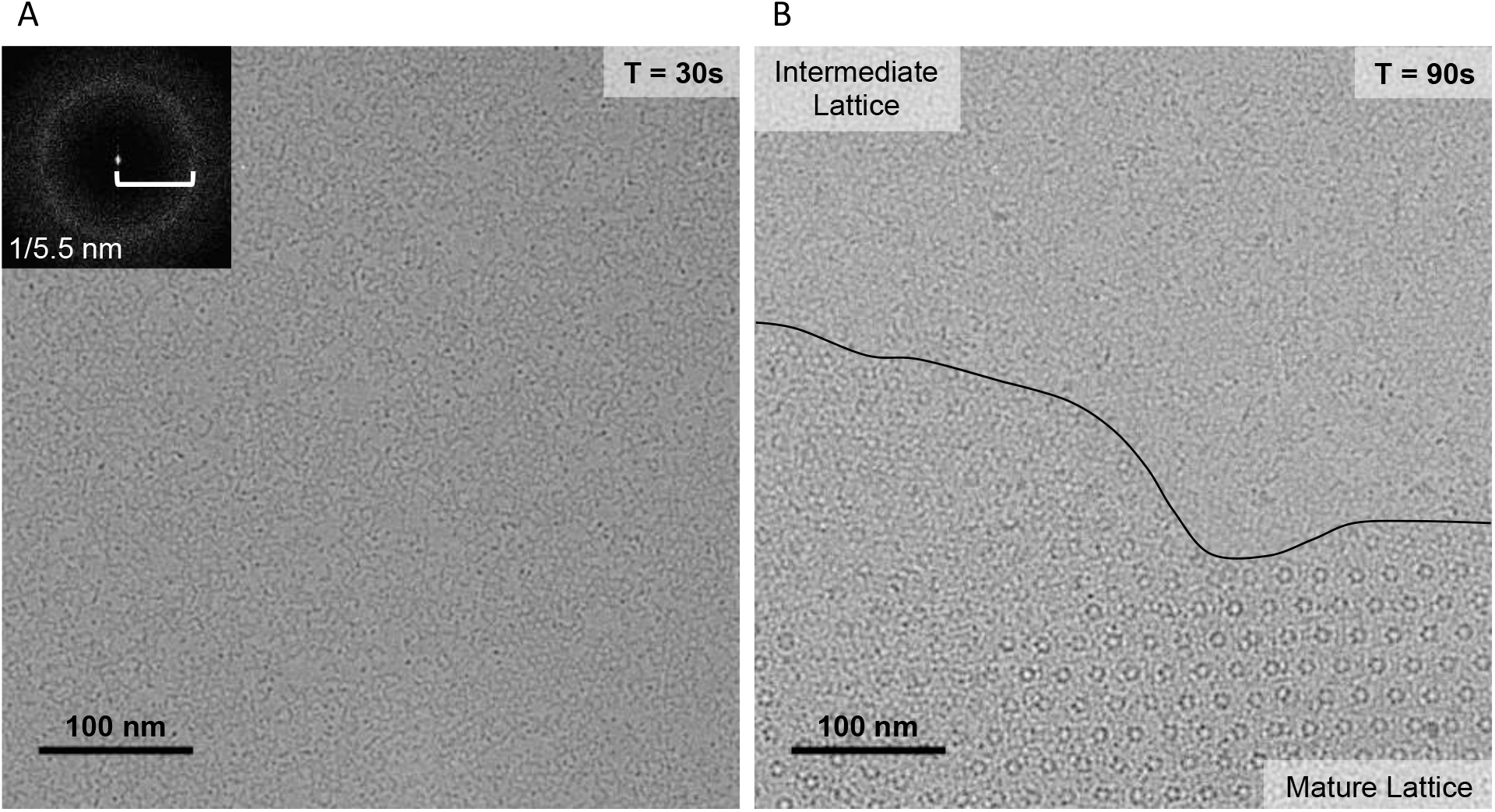
Multiple structural states are observed during RsaA_Full-Length_ self-assembly. A) A Cryo-EM image taken of 20 μM RsaA_Full-Length_ 30 s after adding 10 mM CaCl_2_. The Fourier transform reveals 5.5 nm repeats indicative of the intermediate lattice (inset). B) A Cryo-EM image taken of 20 μM RsaA_Full-Length_ 90 s after adding 10 mM CaCl_2_ reveals the appearance of single and stacked layers of the mature, 22 nm lattice in addition to the intermediate lattice observed in the same image. A black line denotes the boundary between intermediate and mature lattices.

**Figure S7:**
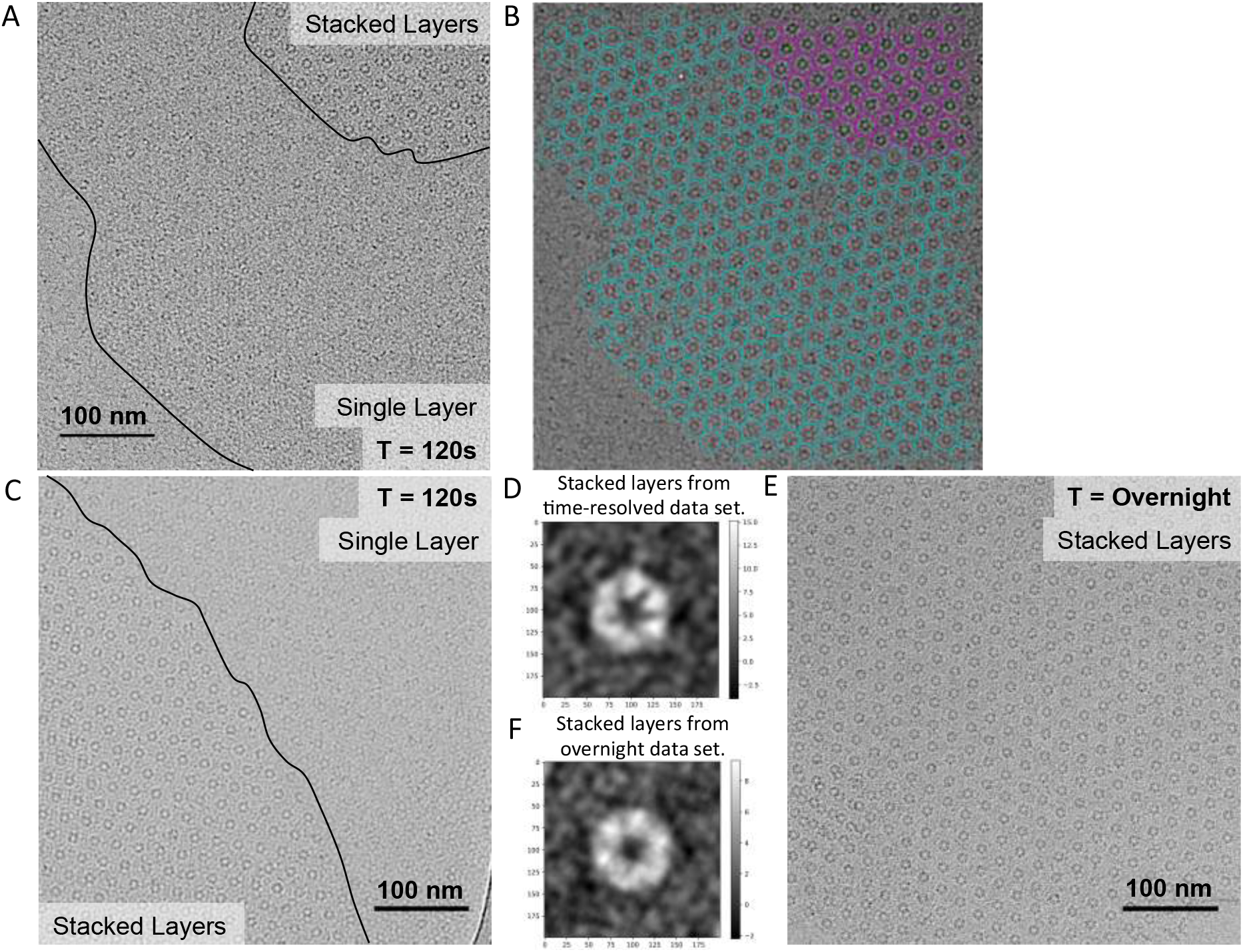
Stacked layers of the RsaA crystal lattice dominate at longer time scales. A) The same Cryo-EM image shown in Figure 3B is reproduced (20 μM RsaA_Full-Length_ 120 s after adding 10 mM CaCl_2_). A single lattice is observed consisting of single layered and stacked layered sections with boundaries denoted by black lines. B) A band-pass filtered version of the image shown in (A) is annotated with stacked layer particles (maroon circles) and single layer particles (cyan circles) identified by Relion. C) Another Cryo-EM image of 20 μM RsaA_Full-Length_ 120 s after adding 10 mM CaCl_2_ is shown. A single lattice consisting of both single and stacked layers is observed with the boundary denoted by a black line. D) A 2D average of 55 stacked layer hexameric particles from the time-resolved data set reveal poor alignment and disappearance of the arms composing the crystallization domain. E) A Cryo-EM image of 20 μM RsaA_Full-Length_ after overnight incubation with 10 mM CaCl_2_ shows only stacked layers. F) A 2D average of 388 stacked layer hexameric particles from crystals grown overnight is similar to that of the time-resolved data set.

**Figure S8:**
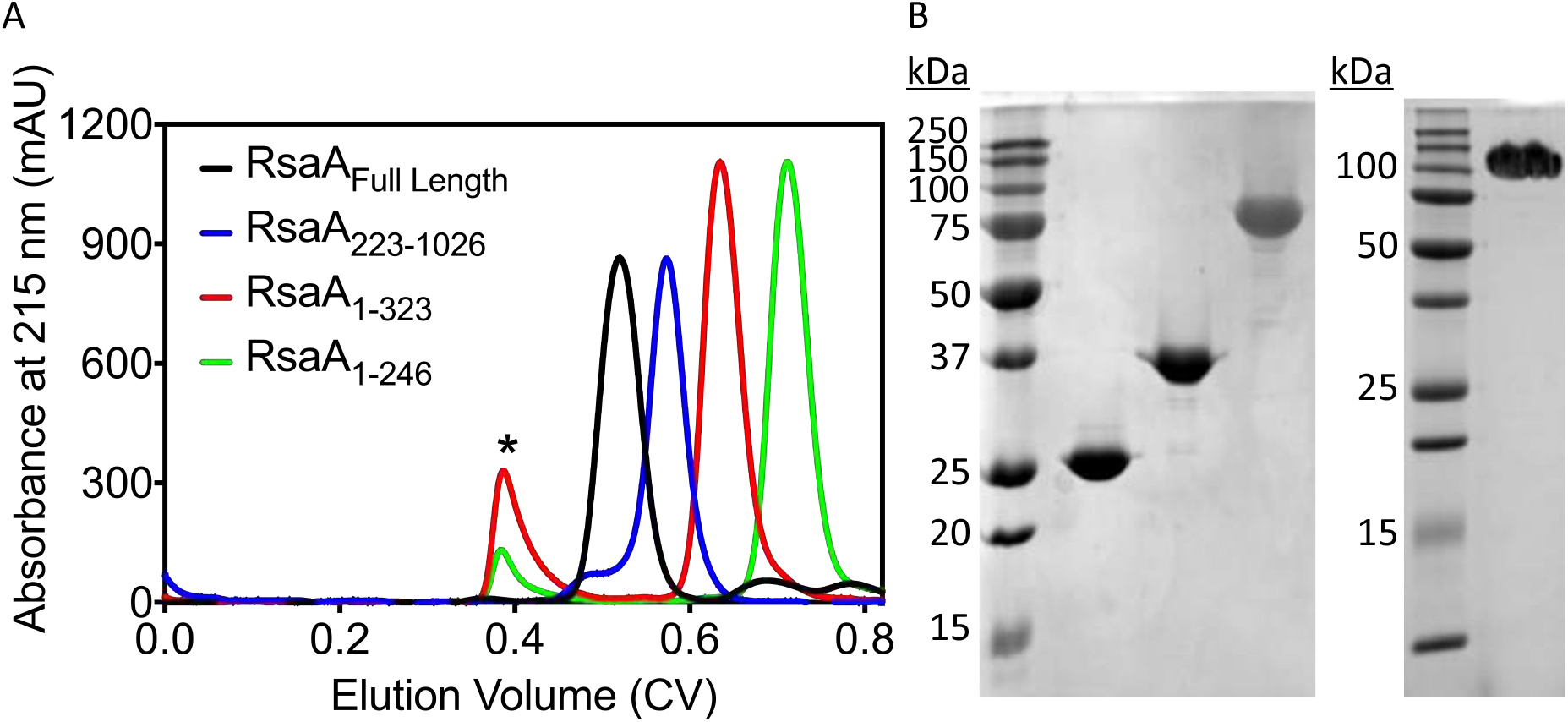
Purification of RsaA_Full-Length_, RsaA_223-1026_, RsaA_1-323_, and RsaA_1-246_. A) Liquid chromatography of RsaA_Full-Length_ (black), RsaA_223-1026_ (blue), RsaA_1-323_ (red), and RsaA_1-246_ (green) using an S200 size exclusion column. A soluble aggregate was observed eluting at the void volume for refolded RsaA_1-323_ and RsaA_1-246_ (asterisk). B) SDS-PAGE of purified samples for RsaA_1-246_, RsaA_1-323_, RsaA_223-1026_, and RsaA_Full-Length_ (left to right).

